# Bacterial sterol methylation confounds eukaryotic biomarker interpretations

**DOI:** 10.1101/2022.05.16.491679

**Authors:** Malory O. Brown, Babatunde O. Olagunju, José-Luis Giner, Paula V. Welander

**Affiliations:** Department of Earth System Science, Stanford University, Stanford, CA 94305; Department of Chemistry, State University of New York-Environmental Science and Forestry, Syracuse, NY 13210

## Abstract

Sterol lipids are required by most eukaryotes and are readily preserved as sterane molecular fossils. These geologic steranes are broadly interpreted as biomarkers for ancient eukaryotes^1,2^ although diverse bacteria also produce sterols^3^. Steranes with side-chain methylations can act as more specific biomarkers^4^ if their sterol precursors are limited to particular extant eukaryotes and are absent in bacteria. An abundance of one such sterane, 24-isopropylcholestane, in late Neoproterozoic rocks has been attributed to marine demosponges and potentially represents the earliest evidence for animals on Earth^5^. However, debates over this interpretation^6–14^ continue given the potential for alternative sources of 24-isopropylcholestane and the lack of experimental evidence demonstrating the function of enzymes that methylate sterols to give the 24-isopropyl side-chain. Here we show that sterol methyltransferases from both sponges and bacteria are functional and identify three bacterial methyltransferases each capable of sequential methylations resulting in the 24-isopropyl sterol side-chain. We identified two of these propylating enzymes in a demosponge metagenome suggesting bacterial symbionts contribute to 24-isopropyl sterol biosynthesis in demosponges. Our results demonstrate yet-uncultured bacteria have the genomic capacity to synthesize side-chain alkylated sterols and should therefore be considered when interpreting side-chain alkylated sterane biomarkers in the rock record.

## Main text

According to the sponge biomarker hypothesis, two side-chain alkylated sterane biomarkers, 24-isopropylcholestane (24-ipc) and 26-methylstigmastane (26-mes), indicate marine sponges of the Demospongiae class^5,15^ (Fig. 1**a**). The co-occurrence of 24-ipc and 26-mes in the late Neoproterozoic (660-540 Ma)^5,15^, preceding the earliest sponge macrofossils by ~100 million years^7^, may represent the first evidence of animals in the geologic record. However, debate over this interpretation continues for several reasons^6–14^ including the potential for alternative sources. Demosponges are the only extant organisms known to contain sterols with the 24-ipc and 26-mes side-chain structures as their major sterols^15–21^, but pelagophytes, which produce 24-*n*-propylcholesterol as their major sterol, also contain 24-isopropyl sterols^22–25^. Trace amounts of 24-ipc and 26-mes were also reported in hydrogenated lipid extracts from Rhizaria^10^, although attempts to replicate these results were not successful^11^. In laboratory pyrolysis experiments under conditions that may mimic diagenesis, 24-ipc and 26-mes form from certain C_29_ sterols suggesting Neoproterozoic 24-ipc and 26-mes could have formed abiotically from algal sterols^13,14^. Further, microbial symbionts, including bacteria from diverse phyla, can constitute up to 40% of sponge biomass^26^. Bacterial symbionts may therefore contribute to 24-isopropyl and 26-methyl sterol biosynthesis in sponges, although bacterial enzymes that methylate the sterol side-chain have not been previously identified.

**Figure 1.**
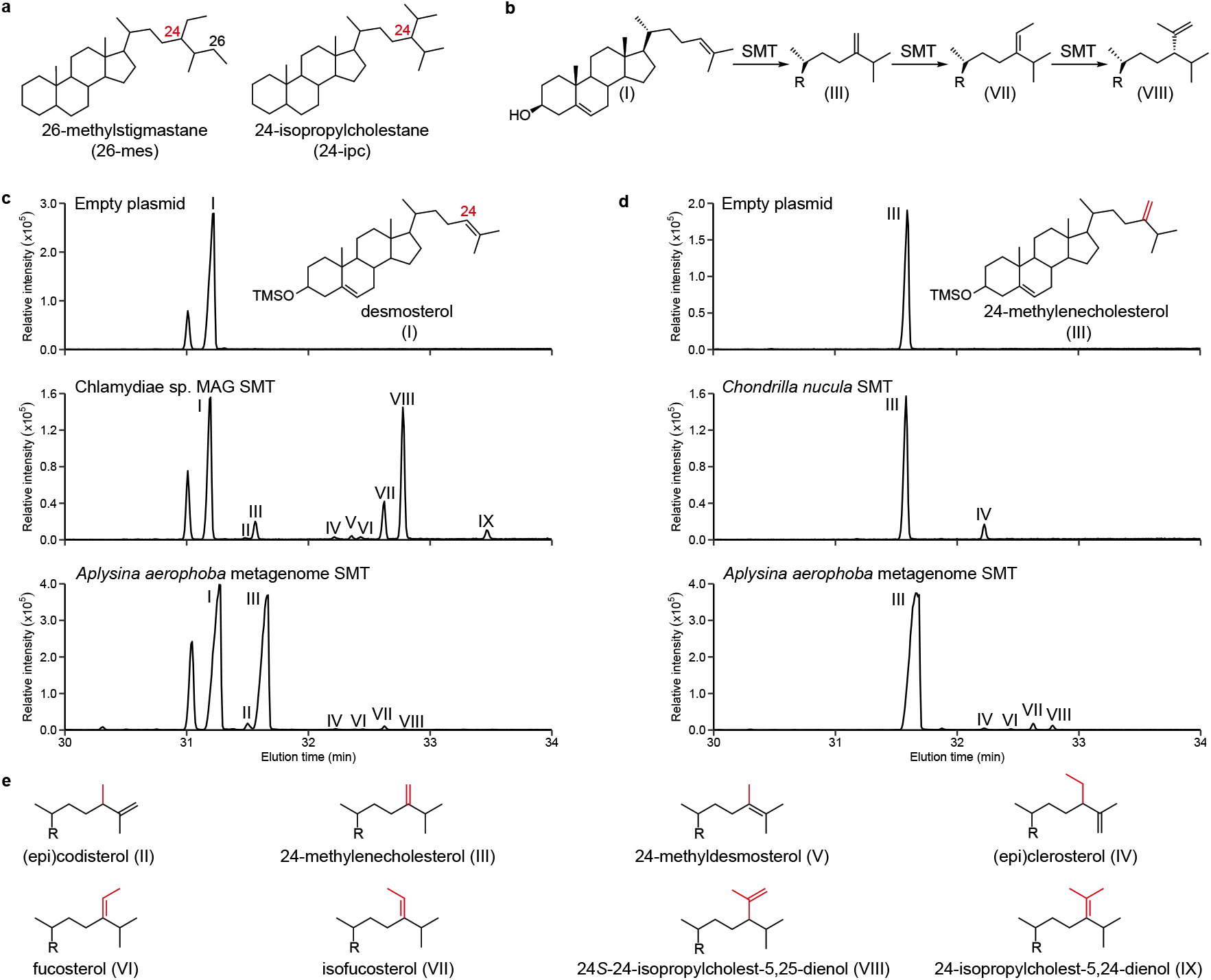
Sterol methyltransferases from sponges and bacteria perform multiple side-chain methylations. **a**, Structures of the sponge biomarkers 26-methylstigmastane (26-mes) and 24-isopropylcholestane (24-ipc). **b**, Predominant pathway to 24-isopropyl sterols by bacteria sterol methyltransferases (SMTs). **c**, Representative extracted ion chromatograms (*m/z* 456, 470, 484, 498) of total lipid extracts from in vitro reactions performed with *E. coli* lysates containing an empty plasmid (**upper**), a sterol methyltransferase (SMT) identified in a freshwater metagenome-assembled genome assigned to the phylum Chlamydiae (**middle**), and an SMT identified in a metagenome from the microbiome of the demosponge *Aplysina aerophoba* (**lower**) with desmosterol as the substrate. **d**, Representative extracted ion chromatograms (*m/z* 456, 470, 484, 498) of total lipid extracts from in vitro reactions performed with *E. coli* lysates containing an empty plasmid (**upper**), an SMT identified in a transcriptome of the demosponge *Chondrilla nucula* (**middle**), and the same SMT identified in the *A. aerophoba* metagenome as in **a** (**lower**) with 24-methylenecholesterol as the substrate. **e**, Side-chain structures of sterols identified in **a** and **b**. All lipids were derivatized to trimethylsilyls prior to GC-MS analysis. Mass spectra of identified sterols are shown in Extended Data Fig. 1.

One approach to constrain uncertainties in the sponge biomarker hypothesis is to identify and characterize the enzymes responsible for these distinct alkylations at C-24 and C-26. Sterol 24-C-methyltransferases (SMTs) alkylate the C-24 position at an isolated double bond in an *S*-adenosylmethionine (SAM) dependent mechanism^27^ and have been characterized extensively in fungi and plants. Bioinformatic analyses addressing this reaction in sponges prompted the hypotheses that at least two SMT copies in a genome are required to produce 24-propyl sterols, and that sponge SMTs can perform multiple methylations at C-24^28^. However, no experimental evidence verifying this activity by any sponge-derived SMT currently exists, and SMTs that produce 24-propyl sterols have not been identified in any extant organism.

### Sponge SMTs are functional

Our first goal was to express putative sponge SMTs and verify methylation activity in vitro. We chose eight SMT homologs identified in genomes and transcriptomes from eight sponges of the Demospongiae, Homoscleromorpha, and Calcarea classes, each of which have been shown to contain 24-ethyl sterols at the species^29–32^ or genus^32–34^ level. We tested each SMT with two potential substrates, desmosterol or 24-methylenecholesterol, and analyzed sterol products using gas chromatography-mass spectrometry (GC-MS). All eight of the sponge SMTs methylated desmosterol to produce 24-methyl sterols including 24-methylenecholesterol and (epi)codisterol (Table 1). Six SMTs also methylated 24-methylenecholesterol to 24-ethyl sterols including fucosterol, isofucosterol, and (epi)clerosterol (Fig. 1**d**). None of the sponge SMTs produced 24-propyl sterols.

**Table 1.**
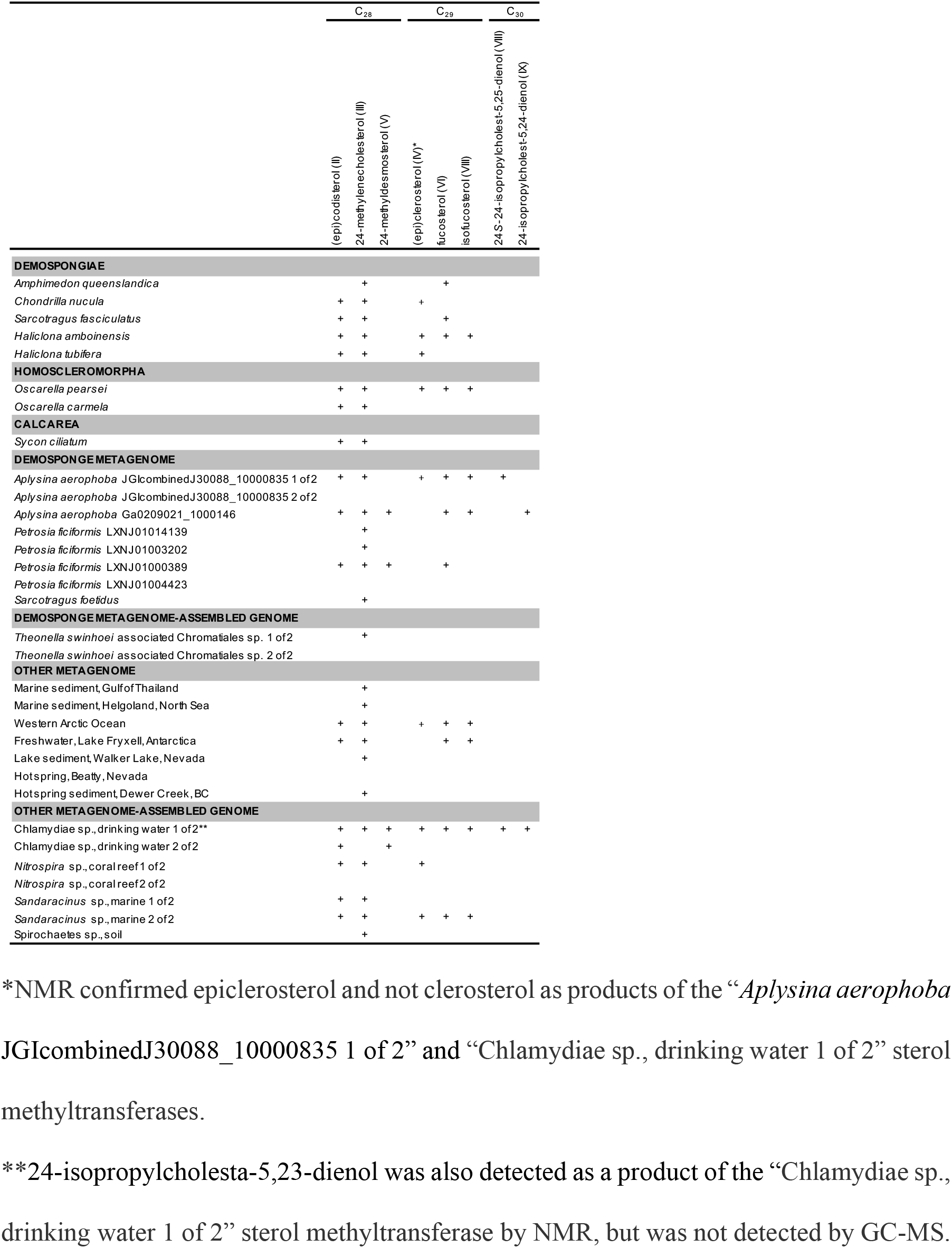
Sterols identified as products of sponge and bacterial sterol methyltransferases.

### Bacterial SMTs produce 24-isopropyl sterols

Our analyses of sponge symbiont metagenomes revealed several SMT homologs, many of which occurred in gene clusters with homologs of other known sterol biosynthesis genes including the bacterial C-4 demethylases *sdmA* and *sdmB^35^* (Fig. 2). We therefore hypothesized bacterial symbionts may contribute to side-chain alkylation in sponges. We chose 10 putative bacterial SMTs identified in metagenomes from four demosponge species for analysis. Eight symbiont SMTs methylated desmosterol to 24-methyl sterols including (epi)codisterol, 24-methylenecholesterol, and 24-methyldesmosterol (Table 1, Fig. 1**c**). Three of these SMTs also methylated 24-methylenecholesterol to 24-ethyl sterols including fucosterol, isofucosterol, and (epi)clerosterol (Fig. 1**d**). Two SMTs identified in a metagenome from *Aplysina aerophoba*, a demosponge known to contain trace 24-isopropyl sterols^21^, also methylated 24-methylenecholesterol to two distinct C_30_ sterols. The first symbiont SMT produced 24*S*-24-isopropylcholest-5,25-dienol while the second produced 24-isopropylcholest-5,24-dienol. Both propylating SMTs were capable of all three methylation steps required to synthesize the 24-ipc side-chain via a single protein. One propylating SMT was identified in a sterol biosynthesis gene cluster suggesting a yet-unknown bacterial sponge symbiont has the genomic capacity to synthesize 24-isopropyl sterols de novo (Fig. 2).

**Figure 2.**
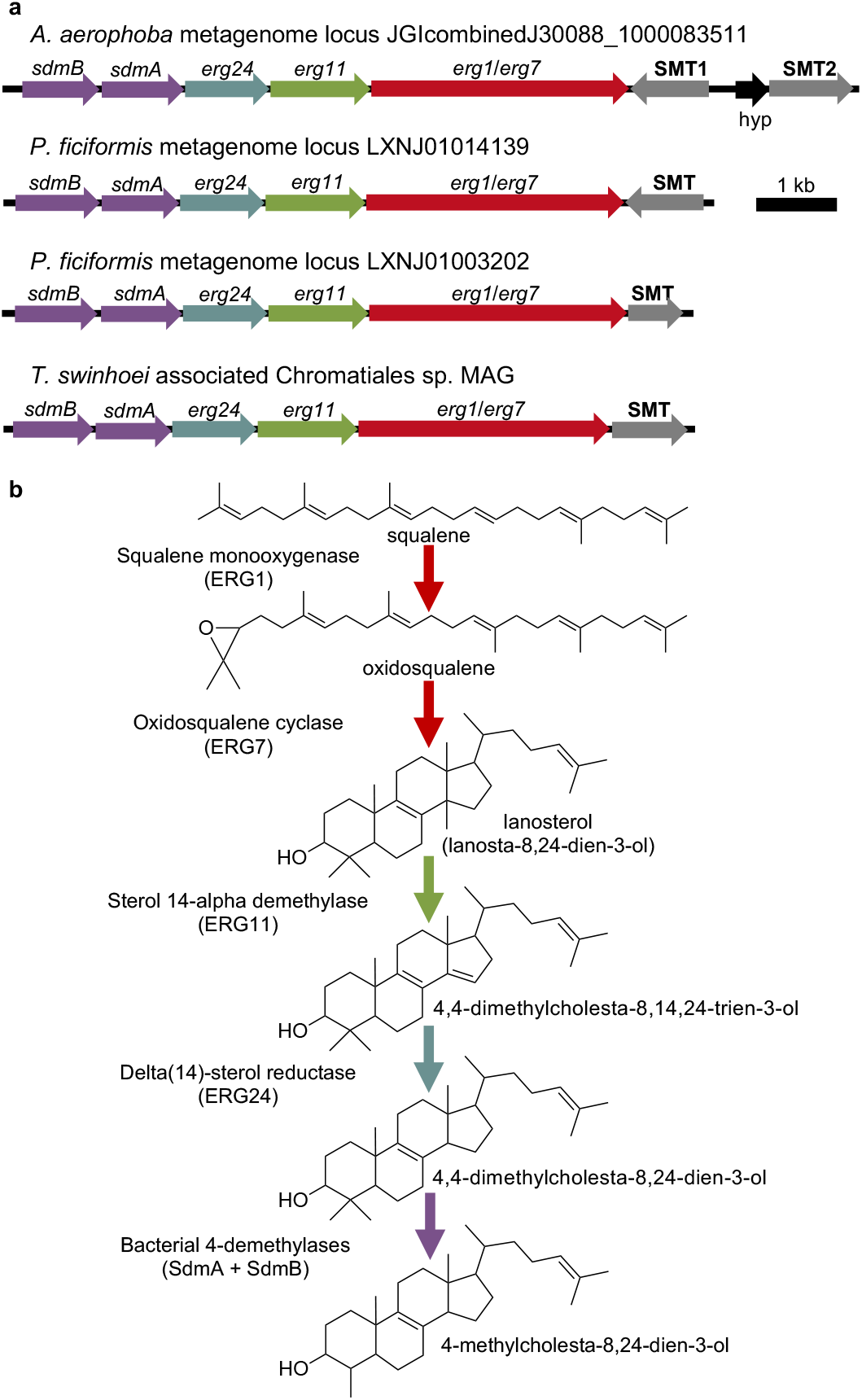
Sterol gene clusters in demosponge metagenomes suggest de novo biosynthesis by bacterial symbionts. **a**, Putative sterol biosynthesis gene clusters identified in metagenomes from the microbiomes of the demosponge species *Aplysina aerophoba, Petrosia ficiformis*, and *Theonella swinhoei*, each containing a functional sterol methyltransferase. **b**, Schematic of the bacterial sterol biosynthesis pathway from squalene to 4-methylcholesta-8,24,-dien-3-ol. Arrow colors at each step correspond to the homologs identifiied in **a**. Detailed information on the source of each metagenomic locus is available in Extended Data Table 1.

Identification of bacterial symbiont SMTs capable of side-chain alkylation led us to question if bacteria from environments outside a sponge host could also produce side-chain alkylated sterols. Through additional searches, we identified SMT homologs in a variety of environmental metagenomes and metagenome-assembled genomes (MAGs). We tested 14 of these SMT homologs from marine, freshwater, thermal spring, and coral reef environments. Twelve environmental SMTs alkylated the sterol side-chain to produce 24-methyl and/or 24-ethyl sterols (Table 1). One SMT, identified in a drinking water filter MAG assigned to the phylum Chlamydiae, produced both 24*S*-24-isopropylcholest-5,25-dienol and 24-isopropylcholest-5,24-dienol (Fig. 1**a**).

The identification of functional metagenomic SMTs likely derived from bacteria, including three that produce 24-isopropyl sterols, is significant given that no bacterial proteins had previously been shown to alkylate the sterol side-chain. To confirm that these SMTs are truly of bacterial origin, we performed phylogenetic analyses of the 24 metagenomic SMTs we tested. In our maximum-likelihood tree of SMT proteins (Fig. 3**a**), the majority of the metagenomic SMTs are distinct and more closely related to each other than to the SMTs of eukaryotes known to produce side-chain alkylated sterols. Further, 23 of the 24 putative bacterial SMTs, including those from sponge metagenomes, form two clades separate from the sponge SMT clade. It is therefore unlikely that these metagenomic SMTs were acquired from sponges via horizontal gene transfer. However, the observed bootstrap values were not robust enough to draw any additional evolutionary conclusions, nor to eliminate any remaining uncertainty of the bacterial origin of these SMTs. We therefore used the recently developed classifier Whokaryote, which predicts whether a metagenomic contig is likely eukaryotic or prokaryotic based on intergenic distance, gene density, gene length, and k-mer frequencies. Whokaryote predicts all 19 of the contigs containing the metagenomic SMTs tested here to be of prokaryotic origin (Extended Data Table 2).

**Figure 3.**
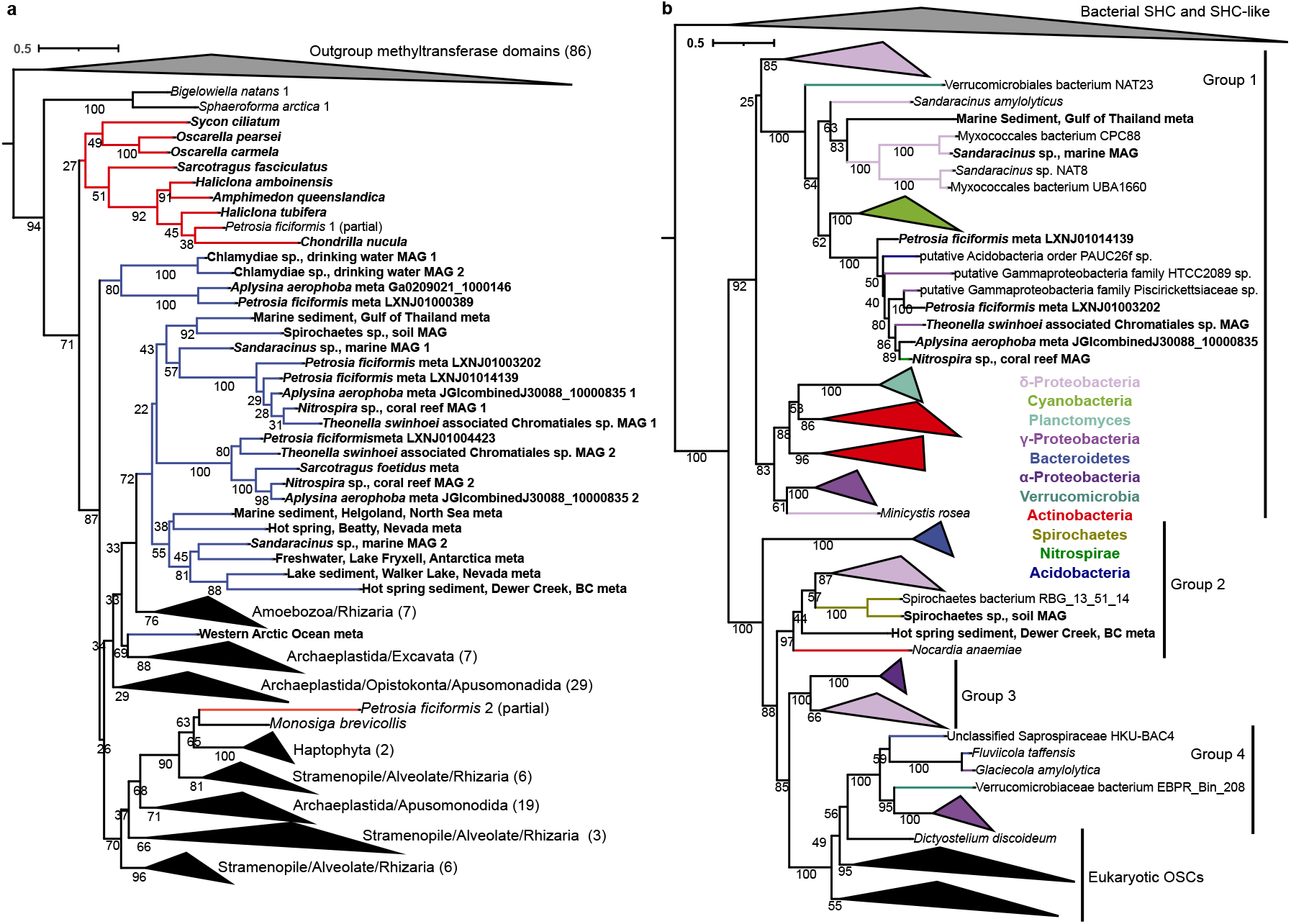
Metagenomic SMTs originate from bacteria. **a**, Maximum likelihood tree generated from a concatenated alignment of the conserved methyltransferase and C-terminal domains of SMT proteins. Metagenomic SMTs are in blue, sponge SMTs are in red, and other eukaryotic SMTs are in black. The numbers in parentheses indicate the number of proteins in collapsed clades. Bolded labels indicate SMTs tested in this study. **b**, Maximum likelihood tree of oxidosqualene cyclases (OSCs). OSC groups are assigned as in^36^. Bolded labels indicate OSCs in metagenomic gene clusters or metagenome-assembled genomes with SMTs tested in this study.

The genomic context of the SMTs we tested provides additional evidence of their bacterial origin. Of the 24 metagenomic SMTs, nine were in gene clusters or MAGs with a homolog of oxidosqualene cyclase (OSC), a key enzyme in sterol biosynthesis responsible for cyclizing oxidosqualene to lanosterol^3^. OSCs cluster into robust phylogenetic clades with high bootstrap support^36^. Thus, phylogenetic analyses of OSCs associated with the SMTs we tested could confirm their bacterial origin. In our maximum-likelihood tree of OSC proteins (Fig. 3**b**), we see that seven of the OSCs associated with the SMTs analysed here, including four from demosponge metagenomes, fall into the Group 1 clade which contains only bacterial OSCs. The remaining 2 OSCs, from soil and hot spring sediment metagenomes, fall into bacterial Group 2. The position of these OSCs provides additional evidence that the metagenomic SMTs we tested originate from a bacterial source.

### Side-chain propylation mechanism

It was surprising to observe that one SMT was capable of propylating the sterol side-chain on its own as it has been hypothesized that at least two SMTs would be required for this reaction^28^. To better understand this unexpected biosynthetic pathway, we performed additional in vitro reactions with two of the bacterial propylating SMTs using ^13^C-labeled sterol substrates and analyzed the resulting products by nuclear magnetic resonance (NMR; Extended Data Table 3). In experiments with [28-^13^C] 24-methylenecholesterol as the substrate of the *A. aerophoba* symbiont SMT, the major C_29_ products were confirmed as isofucosterol and fucosterol present in a 7:1 ratio. Trace epiclerosterol, but not clerosterol, was also detected. The product of triple side-chain methylation was confirmed as *24S*-24-isopropylcholesta-5,25-dienol. The label was found exclusively at C-25, and the *24R* isomer was absent (Extended Data Fig. 2**a**). Fucosterol was not a substrate for the *A. aerophoba* symbiont SMT, but [29-^13^C] isofucosterol yielded *24S*-24-isopropylcholesta-5,25-dienol with the label distributed in a 4:1 label ratio between C-27 and C-26 (Extended Data Fig. 2**b**). This outcome shows interconversion of the C-28 and C-25 cations by hydride-shifts is not involved in the biosynthesis of this compound, and deprotonation of the initially formed cation is unspecific.

The same set of reactions with the propylating Chlamydiae MAG SMT gave similar results. Fucosterol was not a substrate and epiclerosterol, but not clerosterol, was detected. However, minor differences were measured between these two bacterial SMTs. In experiments with [29-^13^C] isofucosterol and the Chlamydiae MAG SMT, the ratio of isofucosterol and fucosterol was 30:1, and the ratio of label at C-27/C-26 of *24S-24-* isopropylcholesta-5,25-dienol was 5:1. Additionally, the C_30_ Δ^23^ and Δ^24^ isomers, 24-isopropylcholesta-5,23-dienol and 24-isopropylcholesta-5,24-dienol, were also detected. The label was detected in the Z-vinylic methyl of 24-isopropylcholesta-5,24-dienol and in one of the two methyls of the Z-isopropyl group of 24-isopropylcholesta-5,23-dienol. These results provide evidence for one hydride-shift to generate the C-24 cation. The ratio of *24S*-24-isopropylcholesta-5,25-dienol and its Δ^24^ and Δ^23^ isomers was estimated to be 20:4:1, respectively. Though minor mechanistic differences between these two propylating SMTs exist, the predominant pathway to 24-isopropyl sterols observed here proceeds from desmosterol to 24-methylenecholesterol to isofucosterol to 24*S*-24-isopropylcholest-5,25-dienol (Fig. 1**b**).

### Discussion

The results presented here experimentally verify that sponges do indeed encode functional SMTs capable of sequentially methylating desmosterol to 24-methylenecholesterol to 24-ethyl sterols, similarly to SMTs recently identified in annelid worms^37^. However, as in Annelida^38^, this activity was inconsistent among Porifera as the SMTs from *Oscarella carmela* (Homoscleromorpha) and *Sycon ciliatum* (Calcarea) did not sequentially methylate sterols, contrary to previous hypotheses^28^. Sponge-derived SMTs that produce 24-ethyl sterols may therefore be restricted to certain species, although we recognize these SMTs may require a different substrate than we provided in our assays. Substrate specificity could also explain why no sponge SMT produced 24-isopropyl sterols in our experiments, although this result was unsurprising given that we did not test SMTs from sponges known to contain propylated sterols due to the lack of available genomic data. However, we did show that bacterial symbionts have the genomic capacity to synthesize these propylated lipids. Previous studies have provided evidence of de novo sterol biosynthesis in demosponges^39–41^, but the experiments were performed prior to the development of 16S community sequencing using cell-free sponge extracts or whole sponge tissue from which the presence of bacterial symbiont-derived enzymes cannot be excluded. Additionally, transcripts for several key sterol biosynthesis genes in the demosponge *Vaceletia* sp. were derived from bacteria^42^. Both sponge and bacterial symbiont proteins may therefore be required to produce 24-isopropyl sterols in vivo. However, the metagenomic gene clusters containing functional SMTs identified here, along with steroid gene clusters containing SMT homologs previously identified in bacterial MAGs associated with demosponges of the *Ircinia* genus^43^, suggest bacterial symbionts may produce 24-isopropyl sterols de novo without demosponge input. Our work highlights the need for robust sequencing of both the sponge host and its microbiome coupled to sterol analysis in demosponge species that contain 24-isopropyls as their major sterols. This may reveal sponge-derived SMTs sufficient for the biosynthesis of these lipids and confirm sponges and/or their bacterial symbionts as potential sources of 24-ipc biomarkers.

The identification of three bacterial SMTs each capable of producing the 24-isopropyl side-chain via a single protein is also significant. Molecular clock analyses suggest demosponges were the likely source of Cryogenian 24-ipc biomarkers given that demosponges acquired a second SMT copy prior to their deposition, while pelagophytes did not acquire a third SMT copy until the Phanerozoic^28^. Additionally, researchers have argued against Rhizaria as a source of 24-isopropyl sterols given that *Bigelowiella natans* encodes only two SMT copies and is thus unlikely to synthesize 24-propyl sterols^11^. These interpretations rely on the hypothesis that multiple SMT copies are required to produce 24-isopropyl sterols. However, our results demonstrating one SMT is sufficient for side-chain propylation make SMT copy number an unreliable predictor of an organism’s ability to produce 24-propyl sterols. Now that propylating SMTs have been identified, site-directed mutagenesis as performed with yeast^44,45^ and plant^46^ SMTs may reveal specific amino acid changes that allow an SMT protein to perform a third methylation. If such key residues are identified, they could act as a more reliable predictor of 24-isopropyl sterol biosynthesis directly from sequencing data than SMT copy number alone.

In addition to its use as a demosponge biomarker, 24-ipc has also been used to indicate marine input into terrestrial systems^47^ given that 24-ipc precursor sterols are only found in marine demosponges and algae. However, the production of 24-isopropyl sterols by an SMT identified in a freshwater Chlamydiae MAG suggests these sterols may occur in freshwater environments. Chlamydiae have not been shown to produce sterols, and homologs of other sterol biosynthesis genes were absent in the Chlamydiae MAG. It is therefore unlikely that this bacterium synthesizes 24-isopropyl sterols de novo. However, some endosymbiotic Chlamydiae recruit sterols from their eukaryotic host into their membranes^48^. Certain freshwater Chlamydiae may therefore methylate host-acquired sterols to give the 24-isopropyl side-chain, which could explain the presence of 22-dehydro-24-isopropylcholestanol throughout a freshwater sediment core from Motte Lake, France^49^. We would therefore exercise caution when interpreting the presence of 24-ipc as evidence for seawater incursion events.

Finally, although we identified SMTs that produce 24-isopropyl sterols, proteins that methylate sterols to give other geologically important side-chain structures remain undiscovered. Identification of enzymes that methylate sterols at the C-26 position to give the 26-mes side-chain structure remains especially important when considering the biomarker evidence for Precambrian sponges. SMTs from sponges and bacteria produced (epi)codisterol suggesting both groups have the genomic capacity to produce substrates for hypothetical 26-SMTs^39,50^. However, it remains to be seen if sponges and bacteria, sponge-associated and otherwise, encode SMTs capable of methylation at C-26. Once identified, in-depth kinetics studies with these enzymes and multiple substrates may reveal if SMTs from sponges and/or bacteria are likely to produce 24-ipc and 26-mes sterol precursors in ratios corresponding to the C_30_ sterane distributions observed in the Cryogenian. Verified side-chain alkylated sterol biosynthesis in a bacterial isolate along with identification of any unique isotopic signatures resulting from this reaction could also confirm bacteria as a source these biomarkers. Until then, the co-occurrence of 24-ipc and 26-mes in high relative abundance continues to provide compelling evidence for derived sponges in the Cryogenian. However, sterol side-chain methylation by symbiotic and free-living bacteria should now be considered when interpreting the presence of ergostane, stigmastane, and 24-isopropylcholestane in the geologic record.

## Methods

### Bioinformatics analysis

SMT homologs were identified in the Joint Genome Institute Integrated Microbial Genomes & Microbiomes (JGI IMG), GenBank, and Compagen (http://www.compagen.org) databases using the *Saccharomyces cerevisiae* S299C SMT amino acid sequence as the BLASTP search query (locus tag: YML008C, maximum e-value: 1e^-50^, minimum percent identity: 30%). Additional demosponge SMT sequences from *C. nucula*, *S. fasciculatas*, and *P. ficiformis* transcriptomes were provided by Prof. David A. Gold^28^. Identifying sequence information is available in Extended Data Table 1. Conserved sterol methyltransferase C-terminal and methyltransferase 11 domains were identified using the HmmerWeb tool (version 2.41.1).

Oxidosqualene cyclase homologs were identified through BLASTP searches of the metagenome scaffolds and MAGs containing the SMTs analyzed in this study and all bacterial and eukaryotic genomes available in JGI IMG as of October 2019 using the *Methylococcus capsulatus* Bath OSC protein sequence (locus tag: MCA2873, maximum e-value: 1e^-50^, minimum percent identity: 20%, minimum sequence length: 350 aa). Redundancy was decreased through manual removal of OSC homologs with 100% similarity at the amino acid level. The N-termini of fused SMO/OSC homologs were trimmed out of the dataset to begin the sequences at the squalene-hopene cyclase N-terminal domain as identified using HmmerWeb.

All protein sequence datasets were aligned via MUSCLE. Conserved SMT domains were aligned separately, then concatenated using Geneious. Maximum-likelihood trees were generated using IQ-Tree on XSEDE with 1,000 ultra-fast bootstrap replicates using the best amino acid substitution models, gamma shape parameters, and invariable sites proportions under the Bayesian Information Criterion and Akaike Information Criterion using ModelTest-NG on XSEDE (Extended Data Table 4). Resulting phylogenetic trees were visualized in iTOL^51^.

All contigs containing metagenomic SMTs were retrieved from JGI IMG or GenBank and analyzed with Whokaryote^52^ using the default inputs with the minimum contig size decreased to 2,000 base pairs. Calculated features and predictions are shown in Extended Data Table 2.

### Gene synthesis and molecular cloning

SMT DNA sequences were codon-optimized for expression in *E. coli* and artificially synthesized by GeneArt (ThermoFisher Scientific) or through the Department of Energy Joint Genome Institute (DOE JGI) DNA Synthesis Science Program. SMT sequences from DOE JGI were obtained in the IPTG-inducible plasmid pSRKGm-*lac*UV5-rbs5^53^ in *E. coli* TOP10. SMT sequences from GeneArt were subcloned into pSRKGm-*lac*UV5-rbs5 by sequence and ligase independent cloning (SLIC)^54^ and transformed into electrocompetent *E. coli* DH10B (Invitrogen) using a MicroPulser Electroporator (BioRad). Oligonucleotides were purchased from Integrated DNA Technologies (Coralville, IA). PCR was performed according the to the manufacturer’s protocol using Phusion DNA Polymerase (New England Biolabs). DNA fragments were purified using the GeneJET Gel Extraction Kit (ThermoFisher Scientific). Plasmid DNA was isolated using the GeneJET Plasmid Miniprep Kit (ThermoFisher Scientific). DNA was sequenced by ELIM Biopharm (Hayward, CA).

### Bacterial culture and heterologous expression

*E. coli* expression strains were cultured in 50 mL terrific broth (TB) supplemented with gentamycin (15 μg/mL) at 37 °C while shaking at 225 rpm. Cultures were induced with 500 μM isopropyl *β*-D-1-thiogalactopyranoside (IPTG) at an OD_600_ of ~0.6, then incubated an additional 4 hours at 30 °C while shaking at 225 rpm. Cells were harvested by centrifugation at 4,500 × *g* for 10 minutes at 4 °C. Cell pellets were stored at −80 °C until sonication.

### Sterol methyltransferase assay

Cells pellets were resuspended in 5 mL buffer containing the following: 50 mM Tris-HCl, 2 mM MgCl_2_, 20% glycerol (v/v), and 0.1% β-mercaptoethanol (v/v), pH 7.5. Cells were then lysed on ice via a Qsonica Sonicator Q500 equipped with a 3.2 mm probe at 30% amplitude pulsing at 5 seconds on, 15 seconds off for 8 minutes of total on time. Lysates were then partially clarified by centrifugation at 4,500 × *g* for 20 minutes at 4 °C. Total protein concentration in the resulting supernatant was quantified using a Coomassie (Bradford) Protein Assay Kit (Thermo Scientific) according to the manufacturer’s Standard Microplate Protocol and a BioTek Synergy HT Microplate Reader. SMT assays were then immediately prepared using enough supernatant to give 4500 μg total protein. Other reaction components included 100 μM desmosterol (Sigma-Aldrich), 24-methylenecholesterol (Avanti Polar Lipids, Inc.), [28-^13^C] 24-methylenecholesterol (Avanti Polar Lipids, Inc.), or synthesized [29-^13^C] fucosterol or isofucosterol, 100 μM *S*-(5’-sdenosyl)-*L*-methionine chloride dihydrochloride (Sigma-Aldrich), and 0.1% Tween-80 (v/v) to a total reaction volume of 400 μL. Reactions were held at 30 °C for 20 hours and then stored at −20 °C until lipid extraction.

### Lipid extraction

Lipids were extracted using a modified Bligh-Dyer method^55,56^ with the completed in vitro reactions as the water phase. Reactions were sonicated in 10:5:4 (vol:vol:vol) methanol:dichloromethane:water for 1 hour. The organic phase was then separated with twice the volume of 1:1 (vol:vol) dichloromethane:water followed by storage at −20 °C for >1 hr. Following centrifugation at 2,800 x *g* for 10 min at 4 °C, the organic phase was transferred and evaporated under N2 to give total lipid extracts (TLEs). TLEs were derivatized to trimethylsilyl ethers in 1:1 (vol:vol) pyridine:Bis(trimethylsilyl)trifluoroacetamide for 1 hr at 70 °C prior to GC-MS analysis.

### GC-MS analysis

Lipids were separated with an Agilent 7890B Series GC equipped with two Agilent DB-17HT columns (30 m x 0.25 mm i.d. x 0.15 μm film thickness) in tandem with helium as the carrier gas at a constant flow of 1.1 ml/min and programmed as follows: 100°C for 2 min, then 12°C/min to 250°C and held for 10 min, then 10°C/min to 330°C and held for 17.5 min. 2 uL of each sample was injected in splitless mode at 250°C. The GC was coupled to an Agilent 5977A Series MSD with the ion source at 230°C and operated at 70 eV in EI mode scanning from 50 to 850 Da in 0.5 s. Sterols were identified based on retention time and comparison to previously published spectra and laboratory standards.

### NMR analysis

In addition to GC-MS analysis, structures of sterol products formed by the metagenomic sterol methyltransferases referred to as “Aplysina aerophoba JGIcombinedJ30088_10000835 1 of 2” and “Chlamydiae sp., drinking water 1 of 2” were confirmed by NMR. ^13^C-labeling experiments were used to assist the identification of the biosynthetic products by 2D-NMR (heteronuclear single quantum coherence (HSQC) and heteronuclear multiple bond correlation (HMBC)) by increasing the amount of ^13^C at a specific position by two orders of magnitude compared to natural abundance. The two terminal isopropyl groups of 24-isopropyl sterols come from biosynthetically disparate origins, the isoprenoid pathway and SAM methylation. Because of the possibilities of hydride-shifts and different positions of deprotonation, each of the four terminal methyl groups can have multiple different origins within the product^20^. In addition to enhanced sensitivity, ^13^C-labeling provided mechanistic details of a pathway which, because of the intrinsic symmetry of 24-isopropyl sterols, would be otherwise hidden.

Total lipid extracts from pooled in vitro reactions using [28-^13^C] 24-methylenecholesterol or [29-^13^C] isofucosterol as the substrate for both SMTs were fractionated using small silica pipet columns (1 mL) eluting with 5 mL each of hexanes:ethyl acetate 9:1, 2:1, and 1:1 (vol:vol). Steroidal constituents were found in the 2:1 solvent fraction and concentrated with a stream of N2 prior to further purification by preparative thin-layer chromatography (TLC; 4:1 hexanes:ethyl acetate). Sterols were separated by high-performance liquid chromatography (HPLC) and the fractions were analyzed by 800 MHz NMR^35^. Sterols were identified by comparison of the HSQC-DEPT and HMBC spectra with reference compounds.

In experiments with [28-^13^C] 24-methylenecholesterol, the presence of isofucosterol was confirmed by a ^1^H-^13^C cross peak corresponding to C-28 at 5.108 and 116.48 ppm. The presence of fucosterol was shown by an HSQC cross peak at 5.183 and 115.58 ppm. Ratios of isofucosterol to fucosterol were determined by comparison of the integral ratios of the cross peaks^35^ for these two isomers. The presence of epiclerosterol was confirmed by cross peaks for C-28 at 1.290/1.380 and 26.01 ppm. Structures of the C29 sterol products were confirmed by long-range HMBC correlations from the 29-methyl group. Isofucosterol showed an HMBC cross peak for H-29 and C-28 at 1.59 and 115.58 ppm; fucosterol and epiclerosterol showed HMBC cross peaks at 1.57 and 116.48 ppm and 0.799 and 26.01 ppm, respectively. The presence of *24S*-24-isopropylcholesta-5,25-dienol labeled at the C-25 position was confirmed by HMBC from the 27-methyl group (cross peak between 1.570 and 147.38 ppm). The presence of 24-isopropylcholesta-5,24-dienol was confirmed by HMBC showing two adjacent cross peaks correlating the C-26 and C-27 methyl protons (1.633/1.651 ppm) with C-25 (123.1 ppm).

In experiments with [29-^13^C] isofucosterol, the label distribution between the C-27 and C-26 positions of *24S*-24-isopropylcholesta-5,25-dienol was shown by integration of the HSQC signals for C-27 (1.570 and 18.95 ppm) and C-30 (4.602/4.740 and 111.85). The label on the Z-vinylic methyl of 24-isopropylcholesta-5,24-dienol was detected by HSQC (1.651 and 19.73 ppm). 24-Isopropylcholesta-5,23-dienol was detected by label at 20.96 ppm exclusively, which is one of the two methyls of the Z-isopropyl group. The ratio of *24S*-24-isopropylcholesta-5,25-dienol, 24-isopropylcholesta-5,24-dienol, and 24-isopropylcholesta-5,23-dienol was estimated by integration of the HSQC.

### Preparation of [29-^13^C] fucosterol and isofucosterol

Specifically labeled C29 sterol precursors were prepared by modification of the procedure reported in^57^. The precursor 24-formylcholesterol i-methyl ether was obtained either through hydroboration of i-Me 24-methylenecholesterol followed by Dess-Martin periodinane oxidation or ozonolysis of i-Me 24-vinylcholesterol. The latter reaction was carried out in 90:9:1 (vol:vol:vol) dichloromethane/methanol/pyridine and yielded the desired 28-formylcholesterol *i*-methyl ether and its methyl hemiacetal in a 1:5 ratio. The hemiacetal was isolated as a mixture of four isomers. ^1^H-NMR (600MHz): 8.06-8.03 (1H, 28-OH), 4.68-4.63 (1H, 28-H), 3.56-3.52 (3H, 28-OMe) 3.332 (s, 3H, 6-OMe), 2.767 (s, 1H, 6-H), 1.108 (s, 3H, 19-H), 0.710 (s, 3H, 18-H). This mixture was reacted with ^13^C-methylmagnesium iodide to give four isomers of [29-^13^C] i-Me 28-hydroxysitosterol: 24R,28R (H-28: 3.938 ppm; C-29: 21.97 ppm), 24S,28S (H-28: 3.938 ppm; C-29: 22.20 ppm), 24R,28S (H-28: 3.820 ppm; C-29: 21.05 ppm), and 24S,28R (H-28: 3.803 ppm; C-29: 21.01 ppm). Preparative TLC (4:1 hexanes:ethyl acetate) gave two bands corresponding to a mixture of the RR and SS isomers (higher Rf) and a mixture of the RS and SR isomers (lower Rf) in ca. 2:1 ratio. These two mixtures were separately dehydrated with phosphorus oxychloride to obtain after deprotection [29 ^13^C] fucosterol (C-29: 13.16 ppm) and [29 ^13^C] isofucosterol (C-29: 12.76 ppm), respectively.

## Data availability

The data that support the findings of this study are available within this paper and its supplementary information files. Additional (raw) data are available from the corresponding author upon reasonable request.

## Acknowledgments

We thank Prof. David A. Gold for providing demosponge SMT protein sequences, Jeremy H. Wei for technical assistance, and Hanon McShea for helpful discussions. We also thank the Department of Energy Joint Genome Institute DNA Synthesis Science Program (Project 503267) for synthesis and cloning of the majority of the SMTs analyzed in this study. Funding for this study was provided by National Science Foundation Grant EAR-1752564 (to P.V.W).

## Author contributions

M.O.B. wrote the manuscript with input from all co-authors. M.O.B. performed the molecular biology experiments, lipid extractions, and GC-MS analyses. M.O.B. and P.V.W. performed the bioinformatic analyses. B.O.O. and J.-L.G. synthesized the ^13^C labeled sterols and performed the NMR analyses. M.O.B analyzed and interpreted the bioinformatic and GC-MS data. B.O.O. and J.-L.G. analyzed and interpreted the NMR data. P.V.W. and M.O.B. conceived the project with contribution from J.-L.G.

---

**Extended Data Figure 1.**
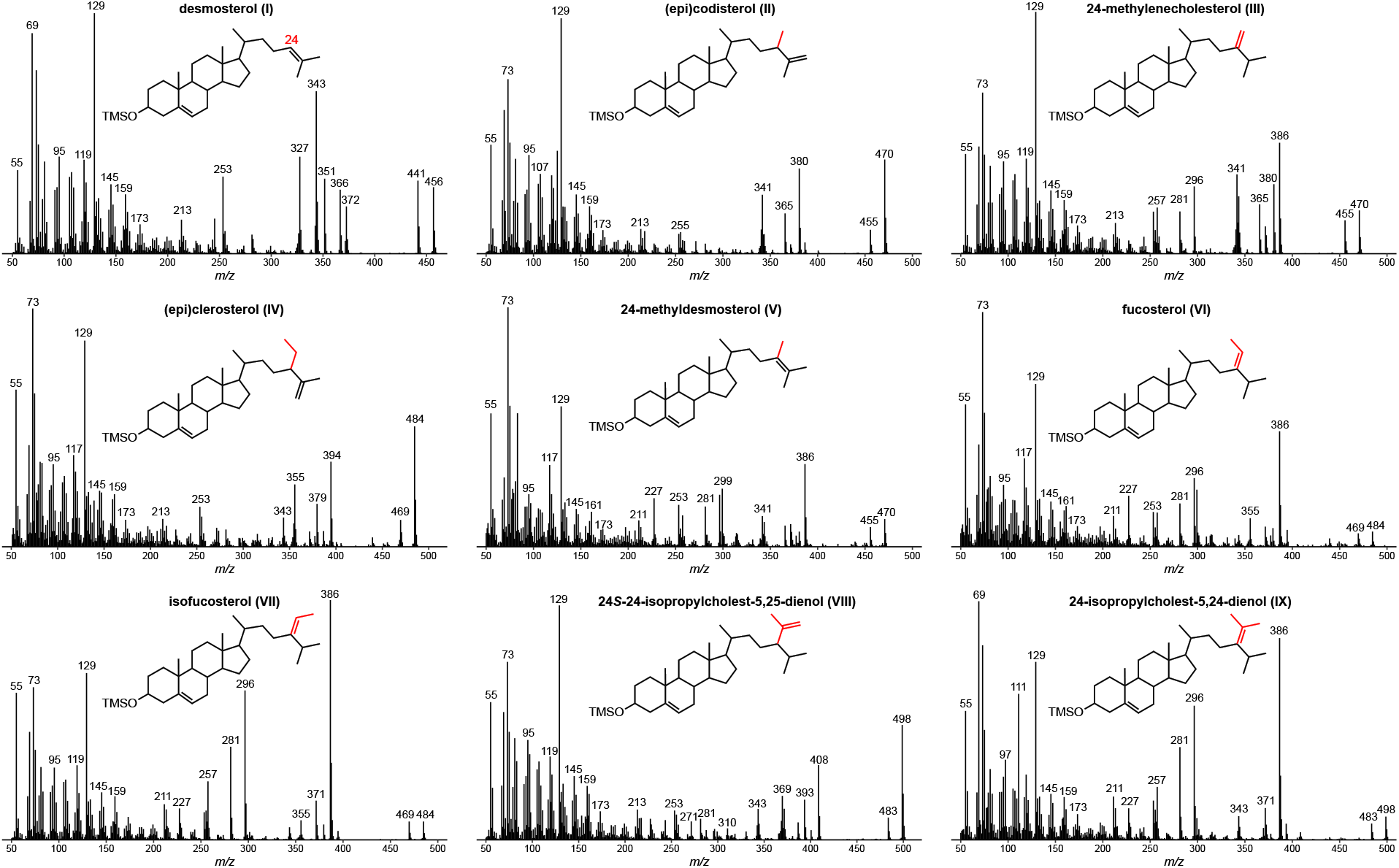
Mass spectra of sterols identified in the main text. Total lipid extracts were derivatized to trimethylsilyl ethers. Detailed GC-MS methods can be found in the Methods section of the main text.

**Extended Data Figure 2.**
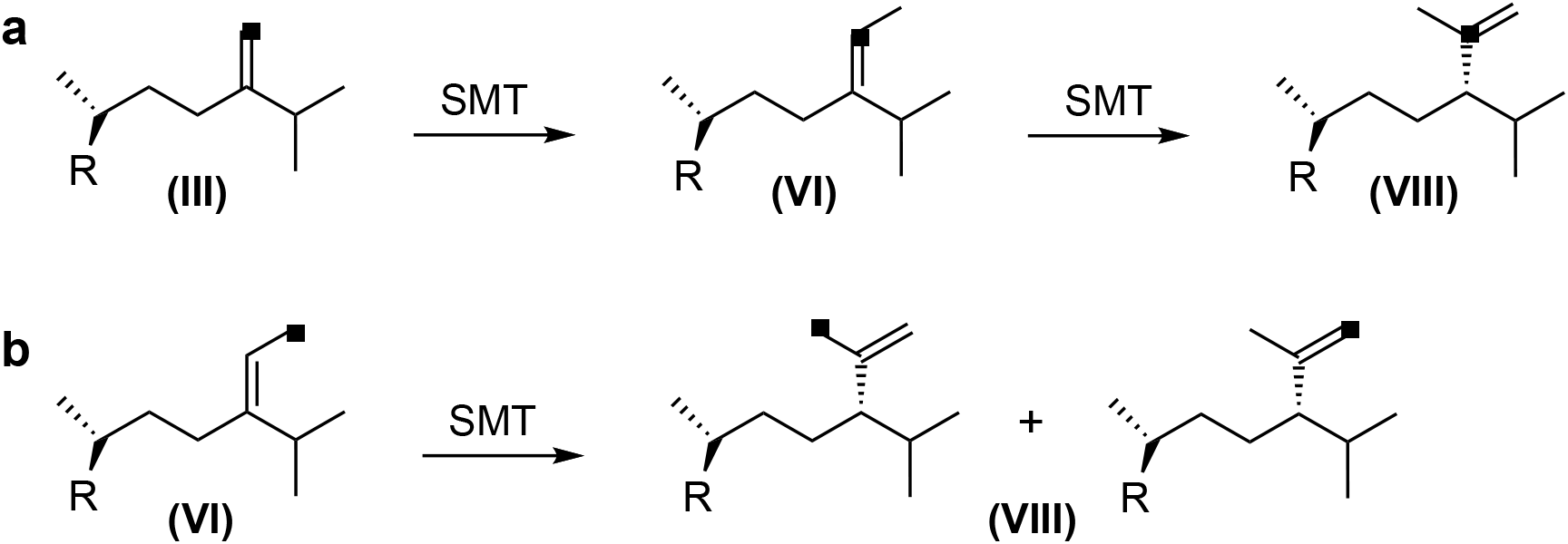
^13^C-labeling experiments with propylating SMTs from an *Aplysina aerophoba* symbiont and a Chlamydiae sp. Positions of the ^13^C labels are shown for the predominant propylation pathway proceeding from 24-methylenecholesterol (III) to isofucosterol (VI) to 24S-24-isopropylcholest-5,25,dienol (VIII). **a**, Experiments with [28-^13^C] 24-methylenecholsterol as the substrate. **b**, Experiments with [29-^13^C] isofucosterol as the substrate.

**Extended Data Table 1.** Sources of sterol methyltransferases analyzed in this study.

| SMT | Source | Genome ID/<br>BioProject | Locus Tag/Locus |
| --- | --- | --- | --- |
| **DEMOSPONGIAE** |  |  |  |
| *Amphimedon queenslandica* | JGI IMG | 2507525020 | anonymous.512 |
| *Chondrilla nucula* | (28) |  |  |
| *Sarcotragus fasciculatus* | (28) |  |  |
| *Haliclona amboinensis* | Compagen |  |  |
| *Haliclona tubifera* | Compagen |  |  |
| **HOMOSCLEROMORPHA** |  |  |  |
| *Oscarella pearsei* | Compagen |  |  |
| *Oscarella carmela* | Compagen |  |  |
| **CALCAREA** |  |  |  |
| *Sycon ciliatum* | Compagen |  |  |
| **DEMOSPONGE METAGENOME** |  |  |  |
| *Aplysina aerophoba* JGIcombinedJ30088_10000835 1 of 2 | JGI IMG | 3300002448 | JGIcombinedJ30088_1000083511 |
| *Aplysina aerophoba* JGIcombinedJ30088_10000835 2 of 2 | JGI IMG | 3300002448 | JGIcombinedJ30088_100008359 |
| *Aplysina aerophoba* Ga0209021_1000146 | JGI IMG | 3300027386 | Ga0209021_100014628 |
| *Petrosia ficiformis* LXNJ01014139 | GenBank | PRJNA318959 | LXNJ01014139.1 |
| *Petrosia ficiformis* LXNJ01003202 | GenBank | PRJNA318959 | LXNJ01003202.1 |
| *Petrosia ficiformis* LXNJ01000389 | GenBank | PRJNA318959 | LXNJ01000389.1 |
| *Petrosia ficiformis* LXNJ01004423 | GenBank | PRJNA318959 | LXNJ01004423.1 |
| *Sarcotragus foetidus* | GenBank | PRJNA318959 | LXNI01004480.1 |
| **DEMOSPONGE METAGENOME-ASSEMBLED GENOME** |  |  |  |
| *Theonella swinhoei* associated Chromatiales sp. 1 of 2 | JGI IMG | 2690315631 | Ga0136241_103361 |
| *Theonella swinhoei* associated Chromatiales sp. 2 of 2 | JGI IMG | 2690315631 | Ga0136241_10845 |
| **OTHER METAGENOME** |  |  |  |
| Marine sediment, Gulf of Thailand | JGI IMG | 3300010430 | Ga0118733_1000126028 |
| Marine sediment, Helgoland, North Sea | JGI IMG | 3300004097 | Ga0055584_1001717352 |
| Western Arctic Ocean | JGI IMG | 3300010883 | Ga0133547_104202862 |
| Freshwater, Lake Fryxell, Antarctica | JGI IMG | 3300009032 | Ga0105048_100346915 |
| Lake sediment, Walker Lake, Nevada | JGI IMG | 3300009504 | Ga0114946_100084975 |
| Hot spring, Beatty, Nevada | JGI IMG | 3300009444 | Ga0114945_100210811 |
| Hot spring sediment, Dewar Creek, BC | JGI IMG | 3300007072 | Ga0073932_10040971 |
| **OTHER METAGENOME-ASSEMBLED GENOME** |  |  |  |
| Chlamydiae sp., drinking water 1 of 2 | JGI IMG | 2619618861 | Ga0074140_11584 |
| Chlamydiae sp., drinking water 2 of 2 | JGI IMG | 2619618861 | Ga0074140_11583 |
| *Nitrospira* sp., coral reef 1 of 2 | JGI IMG | 2681812944 | Ga0131816_12726 |
| *Nitrospira* sp., coral reef 2 of 2 | JGI IMG | 2681812944 | Ga0131816_12722 |
| *Sandaracinus* sp., marine 1 of 2 | JGI IMG | 2775506781 | Ga0248530_10706 |
| *Sandaracinus* sp., marine 2 of 2 | JGI IMG | 2775506781 | Ga0248530_10707 |
| *Spirochaetes* sp., soil | JGI IMG | 2708742746 | Ga0154758_10818 |

**Extended Data Table 2.**
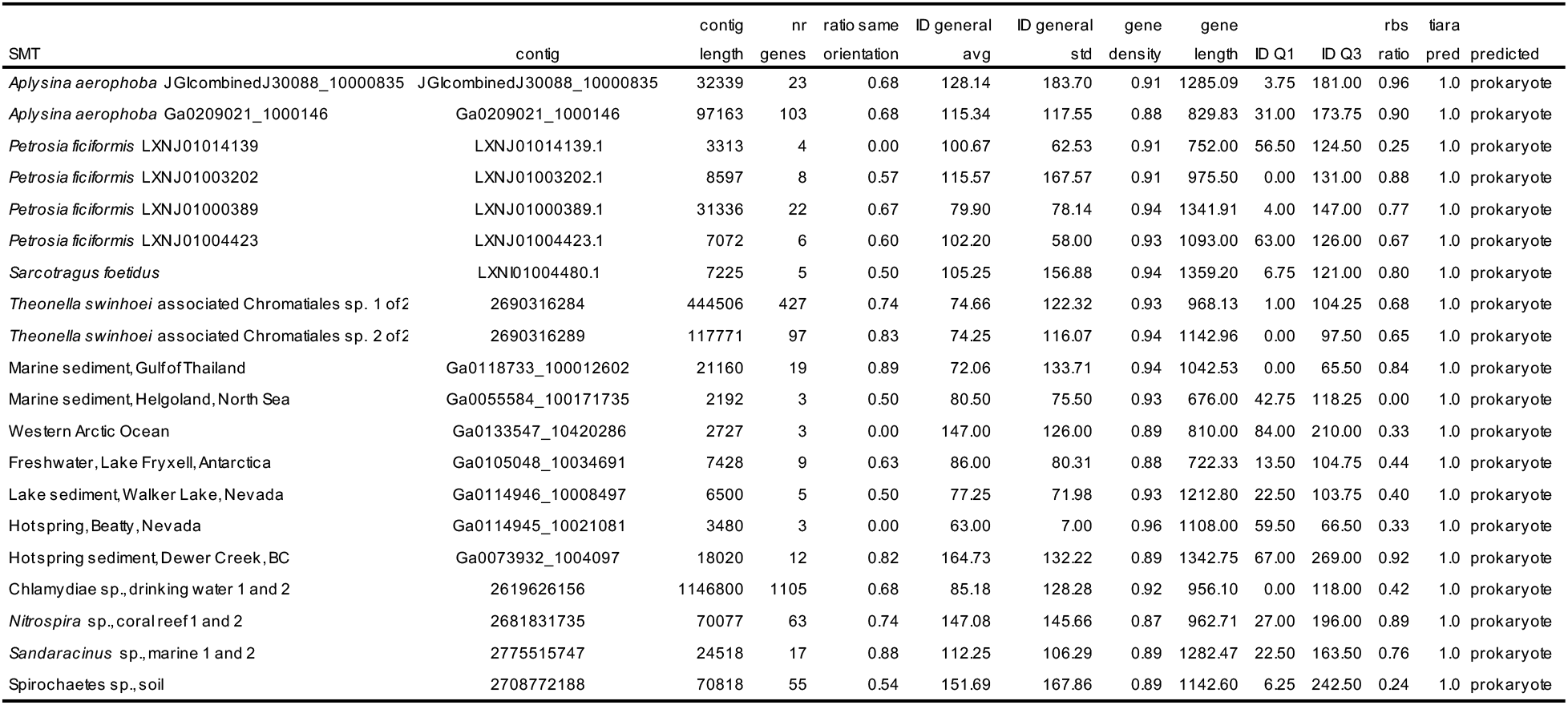
Whokaryote predicts all metagenomic contigs containing sterol methyltransferases analyzed in this study as prokaryotic.

**Extended Data Table 3.** 800 MHz ^1^H and 201 MHz ^13^C NMR assignments of selected sterol side-chain positions at 30 °C.

|  | C-24 | C-25 | C-26 | C-27 | C-28 | C-29 | C-30 |
| --- | --- | --- | --- | --- | --- | --- | --- |
| Fucosterol | **145.6** | 2.199<br>**34.7** |  |  | 5.183<br>**115.58** | 1.574<br>**13.16** |  |
| Isofucosterol | **147.1** | 2.829<br>**28.6** |  |  | 5.108<br>**116.48** | 1.591<br>**12.76** |  |
| Clerosterol* | 1.838<br>**49.54** | **147.61** | 4.730, 4.642<br>**111.33** | 1.570<br>**17.85** | 1.305, 1.341<br>**26.53** | 0.805<br>**12.04** |  |
| Epiclerosterol | 1.81<br>**49.75** | **147.98** | 4.725, 4.636<br>**111.06** | 1.576<br>**18.13** | 1.290, 1.380<br>**26.01** | 0.800<br>**11.96** |  |
| 24*R*-24-Isopropylcholest-5,25-dienol* | 1.56<br>**55.03** | **147.29** | 4.613, 4.738<br>**112.02** | 1.567<br>**18.63** | 1.49<br>**30.25** | 0.911<br>**20.85** | 0.803<br>**21.60** |
| 24*S*-24-Isopropylcholest-5,25-dienol | 1.52<br>**55.50** | **147.38** | 4.602 4.740<br>**111.85** | 1.570<br>**18.95** | 1.51<br>**30.24** | 0.913<br>**20.85** | 0.807<br>**21.49** |
| 24-Isopropylcholest-5,23-dienol |  |  | 1.013<br>**24.77** | 1.013<br>**24.79** |  | 0.980<br>**20.96** | 0.980<br>**21.18** |
| 24-Isopropylcholest-5,24-dienol | **137.0** | **123.1** | 1.633<br>**20.81** | 1.651<br>**19.73** |  | 0.938<br>**21.34** | 0.938<br>**21.41** |

$^{13}\text{C}$ data are in boldface. The assignments were determined by HSQC-DEPT, HMBC, and COSY 2D-NMR experiments. $^1\text{H}$ and $^{13}\text{C}$ chemical shifts are given to three and two decimal places, respectively, if directly observed, two and one, respectively, if determined from 2D spectra. Diastereotopic signals are not assigned. \*Clerosterol and 24*R*-24-isopropylcholest-5,25-dienol were not detected as products of the *A. aerophoba* symbiont and Chlamydiae SMTs but are shown for comparative purposes.

**Extended Data Table 4.** Best amino acid substitution models, gamma shape parameters, and invariable sites proportions under the Bayesian Information Criterion and Akaike Information Criterion using ModelTest-NG on XSEDE for generation of sterol methyltransferase and oxidosqualene cyclase maximum-likelihood trees.

| Protein | Substitution Model | Gamma Shape Parameter | Invariable Sites Proportion |
| --- | --- | --- | --- |
| sterol methyltransferase | LG+I+G | 1.260 | 0.020 |
| oxidosqualene cyclase | LG+I+G | 1.212 | 0.022 |

## Main references

1. Summons, R. E., Bradley, A. S., Jahnke, L. L. & Waldbauer, J. R. Steroids, triterpenoids and molecular oxygen. Philos Trans R Soc Lond B Biol Sci 361, 951–968 (2006).

2. Bobrovskiy, I. et al. Ancient steroids establish the Ediacaran fossil *Dickinsonia* as one of the earliest animals. Science 361, 1246–1249 (2018).

3. Wei, J. H., Yin, X. & Welander, P. V. Sterol synthesis in diverse bacteria. Front Microbiol 7, 990 (2016).

4. Brocks, J. J. et al. The rise of algae in Cryogenian oceans and the emergence of animals. Nature 548, 578–581 (2017).

5. Love, G. D. et al. Fossil steroids record the appearance of Demospongiae during the Cryogenian period. Nature 457, 718–721 (2009).

6. Antcliffe, J. B. Questioning the evidence of organic compounds called sponge biomarkers. Palaeontology 56, 917–925 (2013).

7. Antcliffe, J. B., Callow, R. H. T. & Brasier, M. D. Giving the early fossil record of sponges a squeeze: the early fossil record of sponges. Biol Rev 89, 972–1004 (2014).

8. Love, G. D. & Summons, R. E. The molecular record of Cryogenian sponges - a response to Antcliffe (2013). Palaeontology 58, 1131–1136 (2015).

9. Antcliffe, J. B. The oldest compelling evidence for sponges is still early Cambrian in age - reply to Love and Summons (2015). Palaeontology 58, 1137–1139 (2015).

10. Nettersheim, B. J. et al. Putative sponge biomarkers in unicellular Rhizaria question an early rise of animals. Nat Ecol Evol 3, 577–581 (2019).

11. Love, G. D. et al. Sources of C30 steroid biomarkers in Neoproterozoic–Cambrian rocks and oils. Nat Ecol Evol 4, 34–36 (2020).

12. Hallmann, C. et al. Reply to: Sources of C30 steroid biomarkers in Neoproterozoic–Cambrian rocks and oils. Nat Ecol Evol 4, 37–39 (2020).

13. Bobrovskiy, I. et al. Algal origin of sponge sterane biomarkers negates the oldest evidence for animals in the rock record. Nat Ecol Evol 5, 165–168 (2021).

14. van Maldegem, L. M. et al. Geological alteration of Precambrian steroids mimics early animal signatures. Nat Ecol Evol 5, 169–173 (2021).

15. Zumberge, J. A. et al. Demosponge steroid biomarker 26-methylstigmastane provides evidence for Neoproterozoic animals. Nat Ecol Evol 2, 1709–1714 (2018).

16. Hofheinz, W. & Oesterhelt, G. 24-isopropylcholesterol and 22-dehydro-24-isopropylcholesterol, novel sterols from a sponge. Helv Chim Acta 62, 1307–1309 (1979).

17. Do, H. Q. et al. Unusual sterolic mixture, and 24-isopropylcholesterol, from the sponge *Ciocalypta* sp. reduce cholesterol uptake and basolateral secretion in Caco-2 cells. J Cell Biochem 106, 659–665 (2009).

18. Bortolotto, M., Braekman, J. C., Daloze, D. & Tursch, B. Chemical studies of marine invertebrates. XXXVI. Strongylosterol, a novel C-30 sterol from the sponge *Strongylophora durissima* Dendy. Bull Soc Chim Belges 87, 539–543 (1978).

19. Barnathan, G. et al. Unusual sterol composition and classification of three marine sponge families. Boll Mus Ist Biol Univ Genova 68, 201–208 (2004).

20. Giner, J.-L. Biosynthesis of marine sterol side chains. Chem Rev 93, 1735–1752 (1993).

21. Zumberge, J. A. A lipid biomarker investigation tracking the evolution of the Neoproterozoic marine biosphere and the rise of eukaryotes. (University of California, Riverside, 2019).

22. Kokke, W. C. M. C., Shoolery, J. N., Fenical, W. & Djerassi, C. Biosynthetic studies of marine lipids. 4. Mechanism of side chain alkylation in E-24-propylidenecholesterol by a Chrysophyte alga. J Org Chem 49, 3742–3752 (1984).

23. Giner, J.-L. & Djerassi, C. Biosynthetic studies of marine lipids. 31. Evidence for a protonated cyclopropyl intermediate in the biosynthesis of 24-propylidenecholesterol. J Am Chem Soc 113, 1386–1393 (1991).

24. Giner, J.-L., Li, X. & Boyer, G. L. Sterol composition of *Aureoumbra lagunensis*, the Texas brown tide alga. Phytochemistry 57, 787–789 (2001).

25. Giner, J.-L., Zhao, H., Boyer, G. L., Satchwell, M. F. & Andersen, R. A. Sterol chemotaxonomy of marine pelagophyte algae. C&B 6, 1111–1130 (2009).

26. Taylor, M. W., Radax, R., Steger, D. & Wagner, M. Sponge-associated microorganisms: evolution, ecology, and biotechnological potential. MMBR 71, 295–347 (2007).

27. Nes, W. D. Enzyme mechanisms for sterol C -methylations. Phytochemistry 64, 75–95 (2003).

28. Gold, D. A. et al. Sterol and genomic analyses validate the sponge biomarker hypothesis. Proc Natl Acad Sci U S A 113, 2684–2689 (2016).

29. Gold, D. A. et al. Lipidomics of the sea sponge *Amphimedon queenslandica* and implication for biomarker geochemistry. Geobiology 15, 836–843 (2017).

30. Bergmann, W. & McTigue, F. H. Contributions to the study of marine products. XXI. Chondrillasterol. J Org Chem 13, 738–741 (1948).

31. Venkateswarlu, Y., Reddy, M. V. R. & Rao, M. R. A new epoxy sterol from the sponge *Ircinia fasciculata*. J Nat Prod 59, 876–877 (1996).

32. Fromont, J., Kerr, S., Kerr, R., Riddle, M. & Murphy, P. Chemotaxonomic relationships within, and comparisons between, the orders Haplosclerida and Petrosida (Porifera: Demospongiae) using sterol complements. Biochem Syst Ecol 22, 735–752 (1994).

33. Hagemann, A., Voigt, O., Wörheide, G. & Thiel, V. The sterols of calcareous sponges (Calcarea, Porifera). Chemistry and Physics of Lipids 156, 26–32 (2008).

34. Aiello, A., Fattorusso, E., Magno, S. & Menna, M. Isolation of five new 5α-hydroxy-6-keto-Δ7 sterols from the marine sponge *Oscarella lobularis*. Steroids 56, 337–340 (1991).

35. Lee, A. K. et al. C-4 sterol demethylation enzymes distinguish bacterial and eukaryotic sterol synthesis. Proc Natl Acad Sci USA 115, 5884–5889 (2018).

36. Santana-Molina, C., Rivas-Marin, E., Rojas, A. M. & Devos, D. P. Origin and evolution of polycyclic triterpene synthesis. Mol Biol Evol 37, 1925–1941 (2020).

37. Michellod, D. et al. De novo *phytosterol synthesis in animals*. http://biorxiv.org/lookup/doi/10.1101/2022.04.22.489198 (2022) doi:10.1101/2022.04.22.489198.

38. Gold, D. et al. Sterol methyltransferases in annelid worms rewrite the molecular fossil record. https://www.researchsquare.com/article/rs-1686449/v1 (2022) doi:10.21203/rs.3.rs-1686449/v1.

39. Giner, J.-L. & Djerassi, C. Biosynthetic studies of marine lipids-XXVIII. Use of sponge cell-free extracts in the study of marine sterol biosynthesis. Tetrahedron Lett 31, 5421–5424 (1990).

40. Kerr, R. G., Stoilov, I. L., Thompson, J. E. & Djerassi, C. Biosynthetic studies of marine lipids 16. De novo sterol biosynthesis in sponges. Incorporation and transformation of cycloartenol and lanosterol into unconventional sterols of marine and freshwater sponges. Tetrahedron 45, 1893–1904 (1989).

41. Silva, C. J., Wünsche, L. & Djerassi, C. Biosynthetic studies of marine lipids 35. The demonstration of de novo sterol biosynthesis in sponges using radiolabeled isoprenoid precursors. Comp Biochem Physiol B 99B, 763–773 (1991).

42. Germer, J., Cerveau, N. & Jackson, D. J. The holo-transcriptome of a calcified early branching metazoan. Front Mar Sci 4, 81 (2017).

43. Kelly, J. B., Carlson, D., Low, J. S. & Thacker, R. W. Comparative metagenomics evidence distinct trends of genome evolution between sponge-dwelling bacteria and their pelagic counterparts. http://biorxiv.org/lookup/doi/10.1101/2020.08.31.276493 (2020) doi:10.1101/2020.08.31.276493.

44. Ganapathy, K. et al. Molecular probing of the *Saccharomyces cerevisiae* sterol 24-C methyltransferase reveals multiple amino acid residues involved with C2-transfer activity. Biochim Biophys Acta Mol Cell Biol Lipids 1781, 344–351 (2008).

45. Nes, W. D. et al. Site-directed mutagenesis of the sterol methyl transferase active site from *Saccharomyces cerevisiae* results in formation of novel 24-ethyl sterols. J Org Chem 64, 1535–1542 (1999).

46. Nes, W. D. et al. Probing the sterol binding site of soybean sterol methyltransferase by site-directed mutagenesis: functional analysis of conserved aromatic amino acids in region 1. Arch Biochem Biophys 448, 23–30 (2006).

47. Moldowan, J. M. C30-steranes, novel markers for marine petroleums and sedimentary rocks. Geochim Cosmochim Acta 48, 2767–2768 (1984).

48. Elwell, C. A. & Engel, J. N. Lipid acquisition by intracellular Chlamydiae. Cell Microbiol 14, 1010–1018 (2012).

49. Wünsche, L., Gülaçar, F. O. & Buchs, A. Several unexpected marine sterols in a freshwater sediment. Org Geochem 11, 215–219 (1987).

50. Stoilov, I. L., Thompson, J. E., Cho, J. Ho. & Djerassi, Carl. Biosynthetic studies of marine lipids. 9. Stereochemical aspects and hydrogen migrations in the biosynthesis of the triply alkylated side chain of the sponge sterol strongylosterol. J Am Chem Soc 108, 8235–8241 (1986).

## Methods references

51. Letunic, I. & Bork, P. Interactive Tree Of Life (iTOL) v5: an online tool for phylogenetic tree display and annotation. Nucleic Acids Res 49, W293–W296 (2021).

52. Pronk, L. J. U. & Medema, M. H. Whokaryote: distinguishing eukaryotic and prokaryotic contigs in metagenomes based on gene structure. Microbial Genomics 8, (2022).

53. Banta, A. B., Wei, J. H., Gill, C. C. C., Giner, J.-L. & Welander, P. V. Synthesis of arborane triterpenols by a bacterial oxidosqualene cyclase. Proc Natl Acad Sci USA 114, 245–250 (2017).

54. Li, M. Z. & Elledge, S. J. SLIC: a method for sequence- and ligation-independent cloning. Methods Mol Biol 852, 51–59 (2012).

55. Welander, P. V. et al. Identification and characterization of *Rhodopseudomonas palustris* TIE-1 hopanoid biosynthesis mutants. Geobiology 10, 163–177 (2012).

56. Bligh, E. G. & Dyer, W. J. A rapid method of total lipid extraction and purification. Can J Biochem Physiol 37, 911–917 (1959).

57. Giner, J. L., Zimmerman, M. P. & Djerassi, C. Synthesis of (24R,28R)- and (24S,28S)-24,28-methylene-5-stigmasten-3.beta.-ol and biosynthetic implications of cyclopropyl cleavage to 24-substituted cholesterols. J. Org. Chem. 53, 5895–5902 (1988).

